# Differential contributions of ventral striatum subregions to the motivational and hedonic components of the affective processing of the reward

**DOI:** 10.1101/2021.05.02.442349

**Authors:** Eva R. Pool, David Munoz Tord, Sylvain Delplanque, Yoann Stussi, Donato Cereghetti, Patrik Vuilleumier, David Sander

## Abstract

The ventral striatum is implicated in the affective processing of the reward, which can be divided into a motivational and a hedonic component. Here, we examined whether these two components rely on distinct neural substrates within the ventral striatum in humans. We used a high-resolution fMRI protocol targeting the ventral striatum combined with a Pavlovian-instrumental task and a hedonic reactivity task. Both tasks involved an olfactory reward, thereby allowing us to measure Pavlovian-triggered motivation and sensory pleasure for the same reward within the same participants. Our findings show that different subregions of the ventral striatum are dissociable in their contributions to the motivational and the hedonic component of the affective processing of the reward. Parsing the neural mechanisms and the interplay between Pavlovian incentive processes and hedonic processes might have important implications for understanding compulsive reward-seeking behaviors such as addiction, binge eating, or gambling.

## Introduction

It is widely held that the ventral striatum is implicated in reward processing. The most notorious findings have highlighted the role of the ventral striatum in the computation of reward prediction errors (*1*) and in the anticipation of reward delivery (*2*). Moreover, the ventral striatum has also been consistently implicated in the affective processing of rewarding stimuli (*3, 4*). Affective processes involved in reward processing are often categorized into motivational and hedonic mechanisms. The motivational mechanisms determine how much effort an individual mobilizes to obtain a reward, whereas the hedonic mechanisms determine how much pleasure an individual experiences during reward consumption (*5*).

The ventral striatum itself is not an unitary and homogeneous structure. Studies conducted on rodents and non-human primates typically distinguish between the core and shell nuclei. These nuclei are anatomically distinct, occupying the dorsolateral and ventromedial regions of the ventral striatum. The human ventral striatum is also known to be a heterogeneous structure, but its subdivision into core and shell is not as well defined as it is in rodents and non-human primates. However, recent promising work based on tractographic connectivity profiles suggests that a similar parcellation might also exist in the human ventral striatum (*6–8*). In particular, Cartmell and collaborators (*9*) have provided evidence of a segmentation of the ventral striatum into core-like and shell-like divisions based on a diffusion-tractography analysis of 245 participants. This segmentation of the ventral striatum is very relevant to the affective processing of reward in humans since studies conducted in rodents have consistently demonstrated that the core and shell divisions are differentially involved in the functional processing of motivational and hedonic mechanisms (*5, 10*).

Decades of studies conducted in animals have outlined the critical role of the ventral striatum in reward motivational processes, particularly in Pavlovian-triggered motivation (*10–12*). This type of motivation is usually tested using a key paradigm called Pavlovian-instrumental transfer (PIT). First, individuals learn to associate an instrumental action with a reward, they then learn to associate a Pavlovian stimulus with the reward, and finally, they undergo a transfer test where they are presented with the Pavlovian stimulus in the absence of the reward (i.e., under extinction) and the effort mobilized into the instrumental action is measured. Typically, the Pavlovian stimulus associated with the reward triggers a motivational response enhancing the execution of the instrumental behavior. This phenomenon (known as general Pavlovian instrumental transfer) notably relies on the activity of the core division of the ventral striatum (*10*). Strikingly, animal studies have also demonstrated a critical role of the ventral striatum in hedonic processes (*5, 13*). Hedonic reactivity paradigms measure sensory pleasure in animals by observing their orofacial reactions during food consumption. These hedonic reactions can be amplified by an opioid, orexin, or endocannabinoid stimulation of “hedonic hotspots”. These “hedonic hotspots” are highly focal and distributed across various brain regions such as the insula, the ventral pallidum, and the orbitofrontal cortex (*5, 14*). Critically, “hedonic hotspots” are also found in the shell division of the ventral striatum (*13*). These studies conducted in rodents suggested that within the ventral striatum, there are distinct subregions relying on different neurotransmitters underlying the motivational and the hedonic components of the affective processing of the reward.

A growing number of studies have extended animal findings regarding the role of the ventral striatum in Pavlovian-triggered motivation to humans by adapting the PIT paradigm to a functional Magnetic Resonance Imaging (fMRI) scanner environment (*15–18*). In contrast, the findings suggesting the ventral striatum is involved in sensory pleasure remain less consistent among studies conducted in humans. Similar to animals, sensory pleasure in humans appears to be modulated by opioidergic activity (*19*). However, whereas some studies do find that the magnitude of the experienced sensory pleasure correlates with the activity of the ventral striatum (*20, 21*), sensory pleasure has been most consistently reported to correlate with the activity of the medial orbitofrontal cortex (mOFC) rather than the ventral striatum (*22–24*).

Animal studies suggest that the “hedonic hotspots” in the shell division of the ventral striatum are relatively small (*5*), thereby implying that the standard spatial resolution of fMRI protocols might not be able to reliably detect the hedonic signal from this small region in humans. In recent years, high-resolution fMRI sequences have been developed for the investigation of subcortical regions in reward processing (*25–29*). These protocols record isotropic voxels of 1.8- to 1.5-mm, which are 4 to 5 times smaller than a standard fMRI resolution of 3-mm isotropic voxels. Importantly, these protocols enable researchers to investigate the role of different nuclei in various subcortical structures, thus facilitating the translation of classical animal findings to humans (*26,27*). In the present study, we deployed such a high-resolution protocol to test the hypothesis that distinct subregions of the ventral striatum are differentially involved in processing the Pavlovian motivation and sensory pleasure signals that constitute the affective processing of the reward in humans.

More precisely, we applied a high-resolution fMRI protocol on a human-adapted PIT task (*15, 30*) combined with a hedonic reactivity task; the same olfactory reward was used for both tasks. The PIT task included three phases: an instrumental learning task, a Pavlovian learning task, and a transfer test. During instrumental learning, an instrumental action (i.e., squeezing a handgrip) was first associated with an olfactory reward (unconditioned stimulus, US). Subsequently, during Pavlovian learning, fractal images were either associated with the delivery of the olfactory reward (positive conditioned stimulus; CS+) or with odorless air (negative conditioned stimulus; CS−). The learning of the contingencies between the CSs and the olfactory outcomes was assessed through reaction times in a keypress task and liking ratings of the CSs (*15, 30, 31*). In the final transfer test, the effort mobilized on the handgrip was measured during the presentation of the Pavlovian stimuli (see Fig. 1), the test was administered under extinction, so the olfactory reward was not delivered over this time. The instrumental and Pavlovian learning tasks were administered outside the scanner, whereas the transfer test was administered the following day inside the scanner along with a hedonic reactivity task. In this task, participants smelled the rewarding odor, a neutral odor, and odorless air multiple times. They were asked to report how pleasant the experience of smelling the odor was at that particular time.

**Fig. 1:**
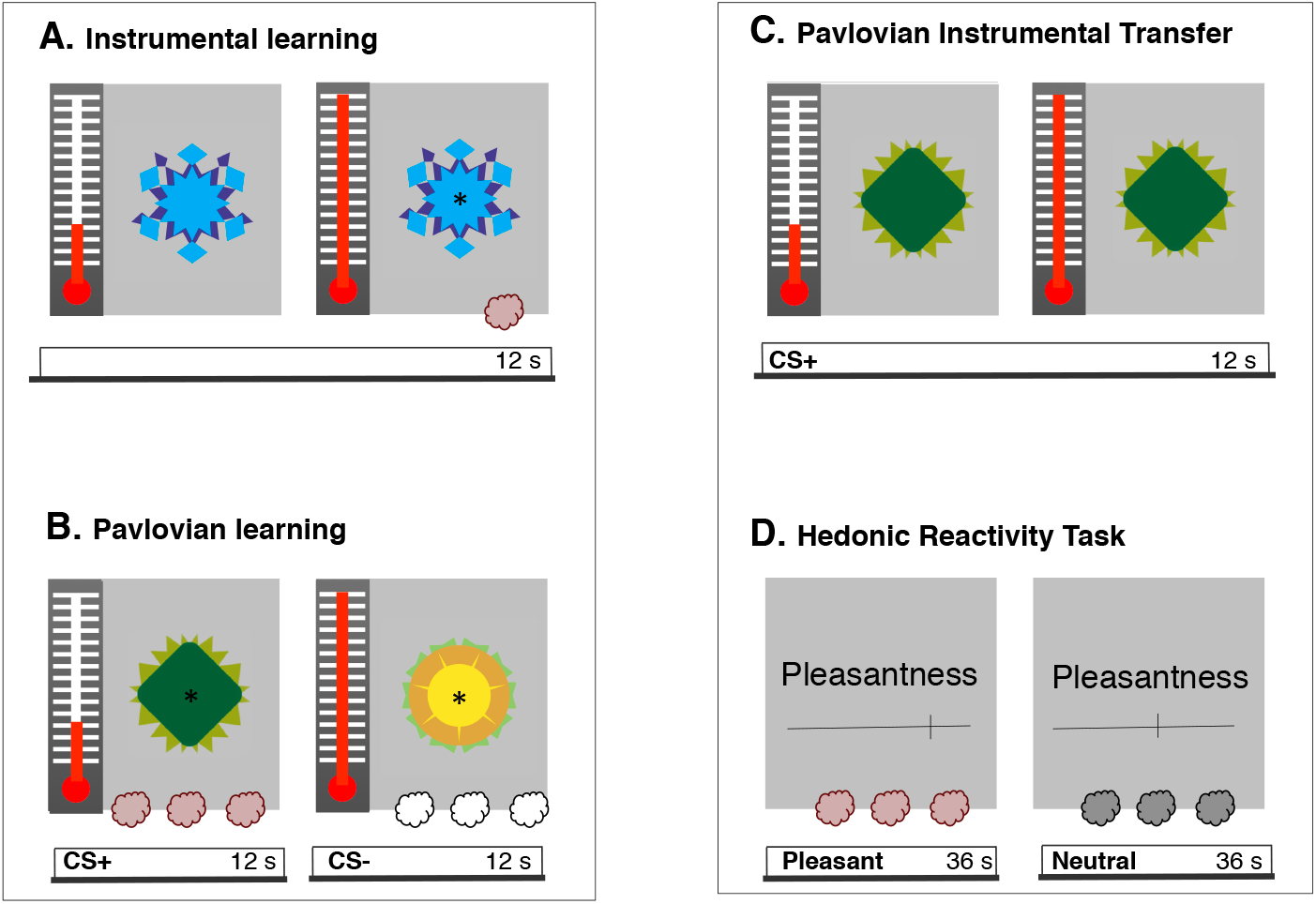
Illustration of the methodological procedure. During day 1 **(A-B)**, participants underwent instrumental and Pavlovian learning outside the scanner. During instrumental learning **(A)**, participants learned to squeeze a handgrip to trigger the release of the olfactory reward. During Pavlovian learning **(B)**, they were exposed to repeated pairings of the positive conditioned stimulus (CS+) with the olfactory reward, whereas the negative conditioned stimulus (CS−) was paired with odorless air. During day 2 **(C-D)**, participants underwent a Pavlovian-instrumental transfer (PIT) test and a hedonic reactivity task inside the scanner. The PIT test **(C)** was administered under extinction. The CS+ and the CS− were displayed in random order (here a CS+ trial is illustrated), and participants could squeeze the handgrip if they wished to do so. The PIT task was adapted from Talmi et al. (*15*). During the hedonic reactivity task **(D)**, participants were presented with the olfactory reward, a neutral odor, and odorless air. They were asked to evaluate on a visual analog scale their perception of the pleasantness (scale from 0 “extremely unpleasant” to 100 “extremely pleasant”) and the intensity (scale from 0 “not perceived” to 100 “extremely strong”) of the odor.

Through this method, we were able to measure separately and without reciprocal contamination the Pavlovian-triggered motivation and the sensory pleasure experience for the same reward within the same participants. Olfactory stimuli have the potential to be powerful reward that have an innate value and biological significance (*32*). They have been successfully used to investigate Pavlovian processes and their underlying neural networks (*31, 33, 34*). Moreover, olfactory rewards present several advantages compared to other kinds of rewards such as money, in that they have the ability to trigger an immediate sensory pleasure experience that can be measured with fMRI. Differently from pictures of food or pictures of money, olfactory stimuli are not a representation of a reward that will be received later, but they are the actual reward that can be consumed immediately, which is an important feature for the empirical measure of hedonic reactions (*35*). Critically, this setting uses experimental tasks that are similar to those used in studies conducted on rodents investigating motivational and hedonic signals within the ventral striatum (*12, 13*). Based on findings in the animal literature, we predicted that the motivational and hedonic components of the affective processing of the reward would rely on distinct subregions of the human ventral striatum.

## Results

### Behavioral results

#### Instrumental conditioning

To test for instrumental learning, we applied a repeated-measures analysis of variance (ANOVA) to the number of squeezes surpassing 50% of each participant’s maximal force (*15,30*) over 24 trials. The analysis did not reveal a statistically significant effect of trial (*F*_(5.08,116.84)_ = 1.54, *p* = 0.181, 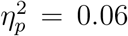, 90% CI = [0.00, 0.11], *BF*_10_ = 0.133; see Fig. 2A). A Post-hoc test revealed very rapid learning, showing that participants significantly increased their responding from the first to the second trial (*F*_(1,23)_ = 24.77, *p* < 0.001, 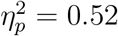, 90% CI = [0.27, 0.68], *BF*_10_ = 312.54; see Fig. 2B).

**Fig. 2:**
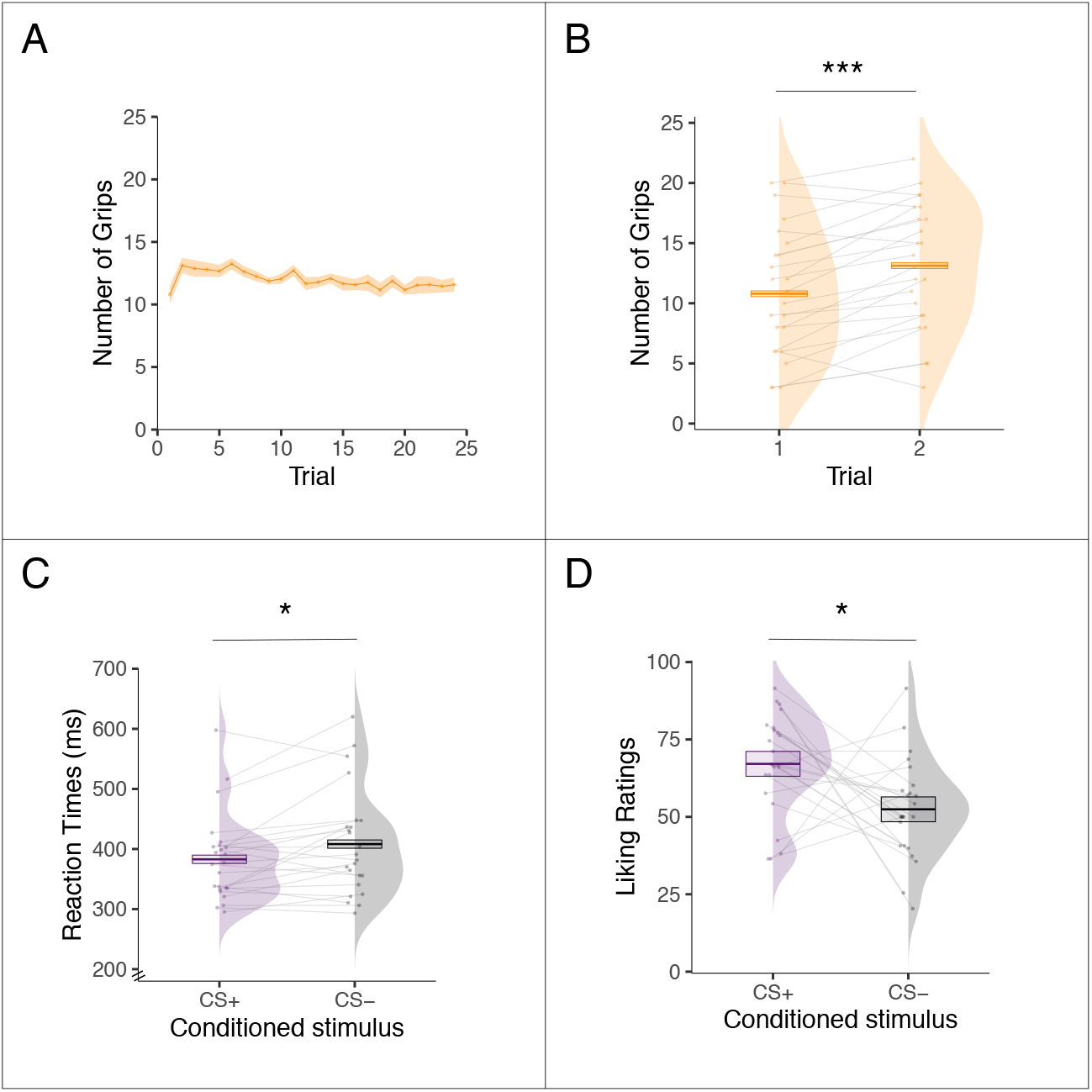
Behavioral results for day 1 outside the scanner. **(A)** Mean number of squeezes on the handgrip during the instrumental learning task displayed as a function of trials over time. **(B)** Mean number of squeezes on the handgrip during the first and second trial of the instrumental learning task. **(C)** Mean reaction times to detect an asterisk while the positive conditioned stimulus (CS+) or the negative conditioned stimulus (CS−) was presented during the Pavlovian learning task. **(D)** Mean liking ratings (scale from 0 “extremely unpleasant” to 100 “extremely pleasant”) of the fractal images used as CS+ and CS− during the Pavlovian learning task. Error bars represent ± 1 *SEM* adjusted for within-participants designs. Asterisks indicate statistically significant differences between conditions (\*\*\**p* < 0.001, \**p* < 0.05)

#### Pavlovian conditioning

To test for Pavlovian learning, we analyzed the reaction times of the keypress task and the liking ratings of the CS images. For the keypress task, we analyzed the reaction times on the first target during the task-on period (*30*). All responses that were more than 3 *SD* from each participant’s mean or absent (2.54% of the trials) were removed. Participants showed evidence of learning in the reaction times: they were faster to detect the target when the CS+ image was presented compared to when the CS− image was presented (*F*_(1,23)_ = 6.67, *p* = 0.017, 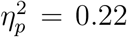, 90% CI = [0.03, 0.45], *BF*_10_ = 3.08; see Fig. 2C). Participants also showed evidence of learning in the liking ratings: they rated the CS+ image as more pleasant than the CS− image (*F*_(1,23)_ = 6.70, *p* = 0.016, 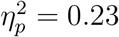, 90% CI = [0.03, 0.45], *BF*_10_ = 18.01; see Fig. 2D).

#### PIT task

We analyzed the number of squeezes surpassing 50% of each participant’s maximal force (*15, 30*) during the transfer test in a 2 (image: CS+ or CS−) by 15 (extinction trials) repeated-measures ANOVA. Participants mobilized more effort when the CS+ image was displayed compared to when the CS− image was displayed (*F*_(1,23)_ = 13.58, *p* < 0.001, 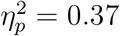, 90% CI = [0.12, 0.57], *BF*_10_ = 29.79; see Fig. 3A,B). There was no statistically significant effect of trial (*F*_(5.07,116.52)_ = 1.39, *p* = 0.23, 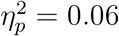, 90% CI = [0.00, 0.10], *BF*_10_ = 0.074) or interaction between trial and CS image (*F*_(5.31,122.12)_ = 0.99, *p* = 0.43, 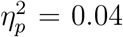, 90% CI = [0.00, 0.07], *BF*_10_ = 0.004; see Fig. 3A).

**Fig. 3:**
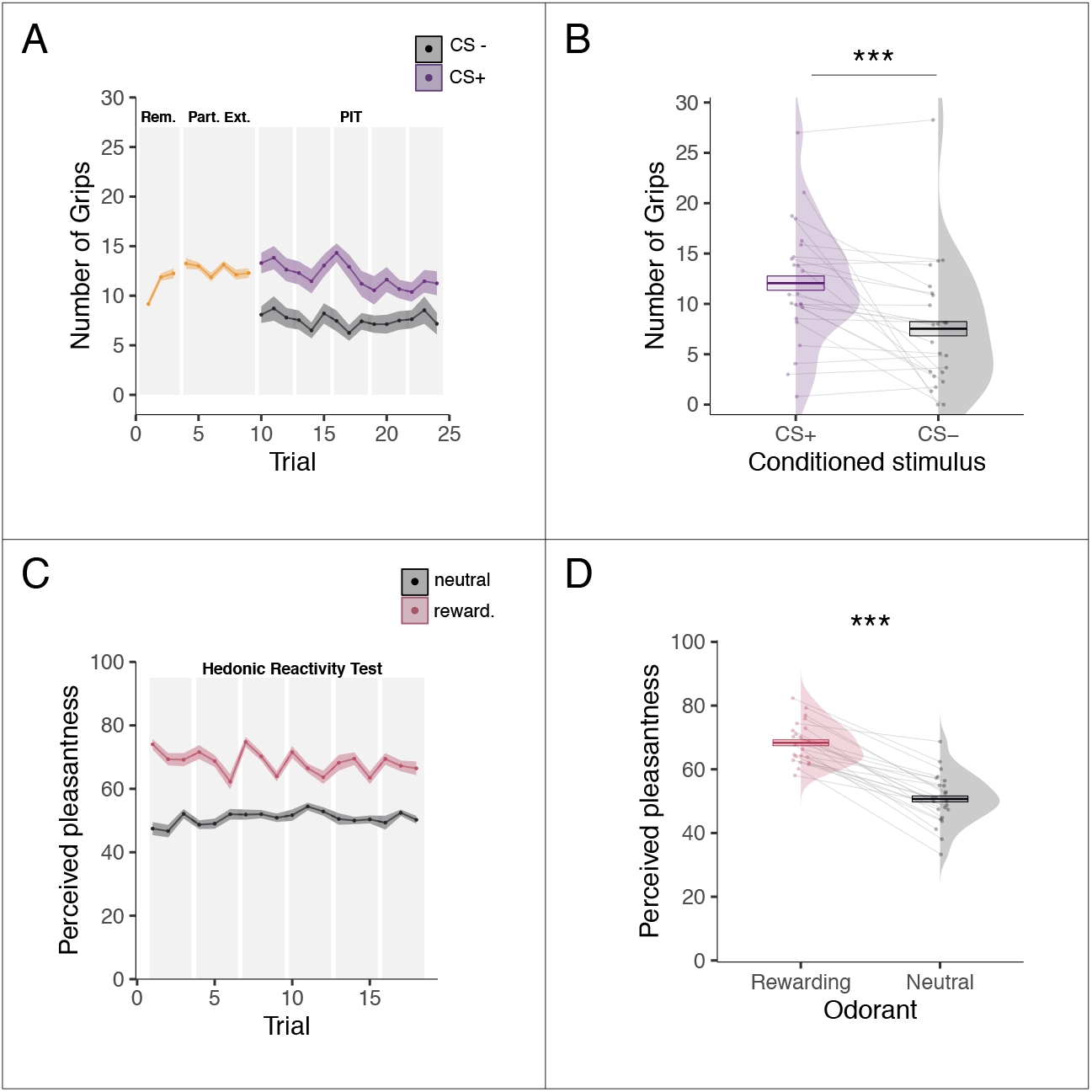
Behavioral results for day 2 inside the scanner. **(A)** Mean number of squeezes as a function of trials over time during the Pavlovian-instrumental transfer (PIT) test. The first 10 trials consisted of a reminder (Rem.) of the instrumental contingencies and a partial extinction (Part. Ext.); the rest of the trials consisted of the actual PIT test for which the number of squeezes is depicted separately for the conditioned stimulus previously associated with the olfactory reward (CS+) and the conditioned stimulus previously paired with odorless air (CS−). **(B)** Overall mean number of squeezes while the CS+ or the CS− was presented during the PIT test. **(C)** Mean perceived pleasantness ratings as function of trials over time depicted separately for the rewarding and the neutral odors during the hedonic reactivity task. **(D)** Overall mean perceived pleasantness of the rewarding and the neutral odors. Pleasantness was evaluated on a scale going from 0 (”extremely unpleasant”) to 100 (”extremely pleasant”) Error bars represent ± 1 *SEM* adjusted for within-participants designs. Asterisks indicate statistically significant differences between conditions (\*\*\**p* < 0.001)

#### Hedonic reactivity task

We analyzed the pleasantness ratings during the hedonic reactivity task with a 2 (odor: rewarding or neutral) by 18 (trial) repeated-measures ANOVA. As expected, participants rated the olfactory reward as more pleasant than the neutral odor (*F*_(1,23)_ = 136.66, *p* < 0.001, 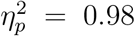, 90% CI = [0.97, 0.99], *BF*_10_ = 4.1 × 10^9^; see Fig. 3B,C). The analysis additionally showed a main effect of trial (*F*_(8.42,193.55)_ = 2.19, *p* = 0.028, 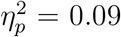, 90% CI = [0.01, 0.12], *BF*_10_ = 3.06; see Fig. 3C) and an interaction between odor and trial (*F*_(8.94,205.52)_ = 4.29, *p* < 0.001, 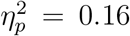, 90% CI = [0.06, 0.21], *BF*_10_ = 5863.90; see Fig. 3C). A follow-up analysis showed that the perceived pleasantness of an odor at trial t was influenced by the value of the preceding odor at trial t-1: The olfactory reward was rated as more pleasant when it was preceded by the neutral odor compared to when it was preceded by another olfactory reward (*F*_(1.35,31.08_ = 97.43, *p* < 0.001, 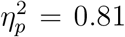, 90% CI = [0.70, 0.87], *BF*_10_ = 3.01 × 10^15^).

Because the neutral and rewarding odors were selected during day 1 (outside the scanner) to have similar intensities, we also analyzed the intensity ratings during the hedonic reactivity task as a control. Participants rated the olfactory reward as more intense than the neutral odor (*F*_(1,23)_ = 15.87, *p* < 0.001, 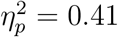, 90% CI = [0.15, 0.60], *BF*_10_ = 73.96). There was also a main effect of trial (*F*_(7.90,181.80)_ = 9.25, *p* < 0.001, 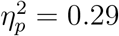, 90% CI = [0.18, 0.35], *BF*_10_ = 3.06), but no statistically significant interaction between odor and trial emerged (*F*_(8.46,194.61)_ = 0.94, *p* = 0.49, 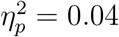, 90% CI = [0.00, 0.05], *BF*_10_ = 0.002). A follow-up analysis showed that the odor at trial t was perceived as more intense when it was preceded by a different odor at trial t-1 compared to when it was preceded by the same odor at trial t-1 (*F*_(1,23_ = 57.74, *p* < 0.001, 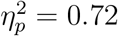, 90% CI = [0.53, 0.81], *BF*_10_ = 4060.65).

### fMRI results

#### Tasks validation

Before focusing on our hypotheses in our region of interest, we validated our paradigms and the quality of our signal through two control analyses. We report the results from our analyses within predefined ROI in the olfactory cortex, the cerebellum, and the thalamus using a height threshold of *p* < 0.005, with an extent threshold significant at *p* < 0.05 corrected for multiple comparisons. For the hedonic reactivity task, the odor presence (odor > odorless air) activated the piriform bilaterally (*k*^*thr*^ = 18; right: MNI [*xyz*] = [25 −3 −20], *k* = 89, *β* = 0.54, 95% CI = [0.37, 0.70], *SE* = 0.079; left: MNI [*xyz*] = [−23 −7 −10], *k* = 47, *β* = 0.52, 95% CI = [0.32, 0.71], *SE* = 0.096; see Fig. 4A). For the PIT task, the overall frequency of the squeezes executed with the right hand activated the motor regions in our field of view. More precisely, the right cerebellar hemisphere (*k*^*thr*^ = 44; MNI [*xyz*] = [18 −48 −18], *k* = 1128, *β* = 1.04, 95% CI = [0.70, 1.38], *SE* = 0.164; see Fig 4B), and left thalamus, (*k*^*thr*^ = 37; MNI [*xyz*] = [−16 −20 7], *k* = 168, *β* = 0.45, 95% CI = [0.32, 0.59], *SE* = 0.065; see Fig. 4B).

**Fig. 4:**
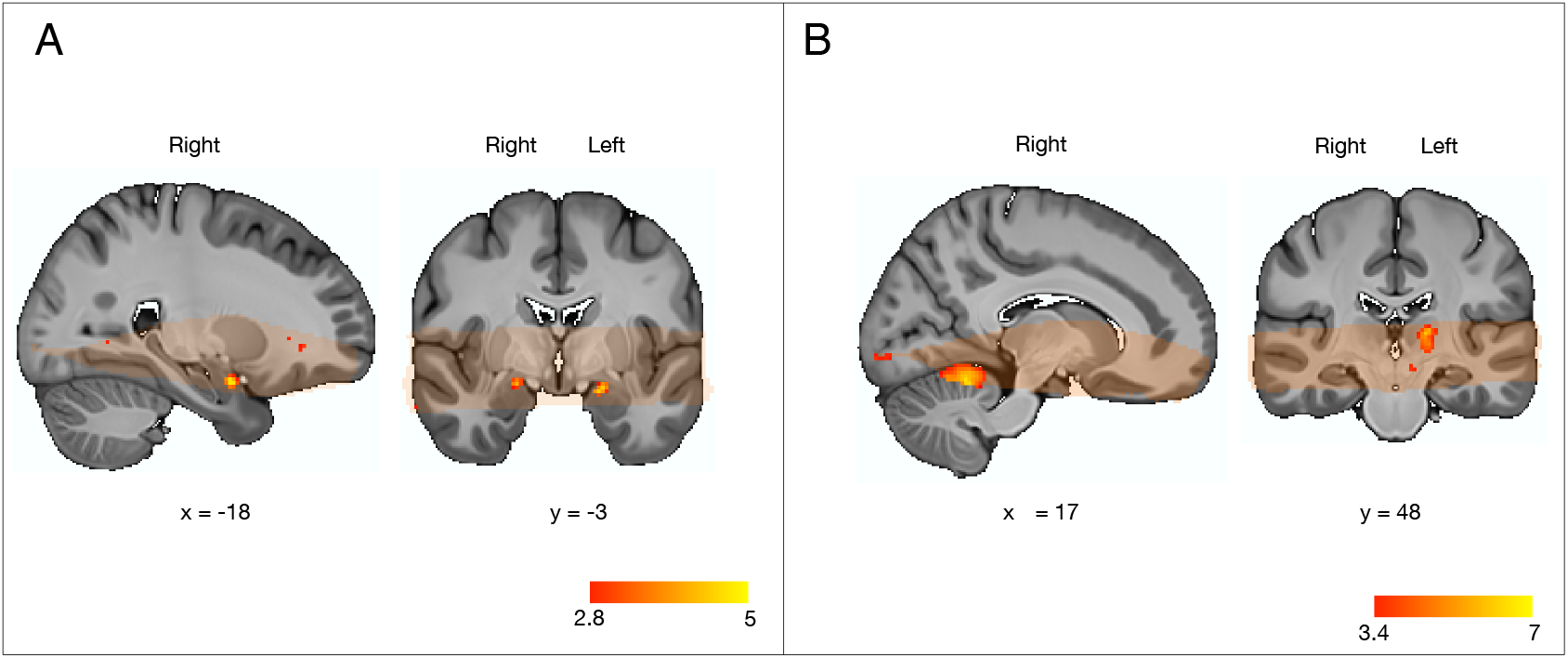
Olfactory and motor s ignals in the hedonic reactivity task and the PIT task. **(A)** An olfactory signal was found in the bilateral piriform cortex during the hedonic reactivity task. For display purposes statistical *t*-maps are shown with a threshold at *p* < 0.005, uncorrected **(B)**. A motor signal was found in the right cerebellum and the left thalamus during the PIT task. For display purposes, statistical *t*-maps are shown with a threshold at *p* < 0.001, uncorrected. Orange overlays indicate brain areas from which functional MRI data was acquired in all participants and were thus included in the statistical analysis. Scale bar shows *t*-statistic.

#### PIT task

We report the results from our analyses within the predefined ROI in ventral striatum and mOFC using a height threshold of *p* < 0.005, with an extent threshold significant at *p* < 0.05 corrected for multiple comparisons.

Following the between-participants analysis typically used in PIT tasks (*15, 27*), we extracted the CS+ vs. CS− contrast for each participant at the first-level and correlated it with the average PIT effect (increased effort during the CS+ compared to the CS−) of each participant at the second-level.

For this contrast, we did not find any significant activation in the mOFC, but as shown in Fig. 5A and B we found a bilateral activation of the dorsolateral subregion of the ventral striatum (*k*^*thr*^ = 16; left: MNI [*xyz*] = [−18 23 −4], *k* = 69, *β* = 0.11, 95% CI = [0.080, 0.15], *SE* = 0.017; right: MNI [*xyz*] = [13 13 −2], *k* = 159, *β* = 0.12, 95% CI = [0.085, 0.16], *SE* = 0.017).

**Fig. 5:**
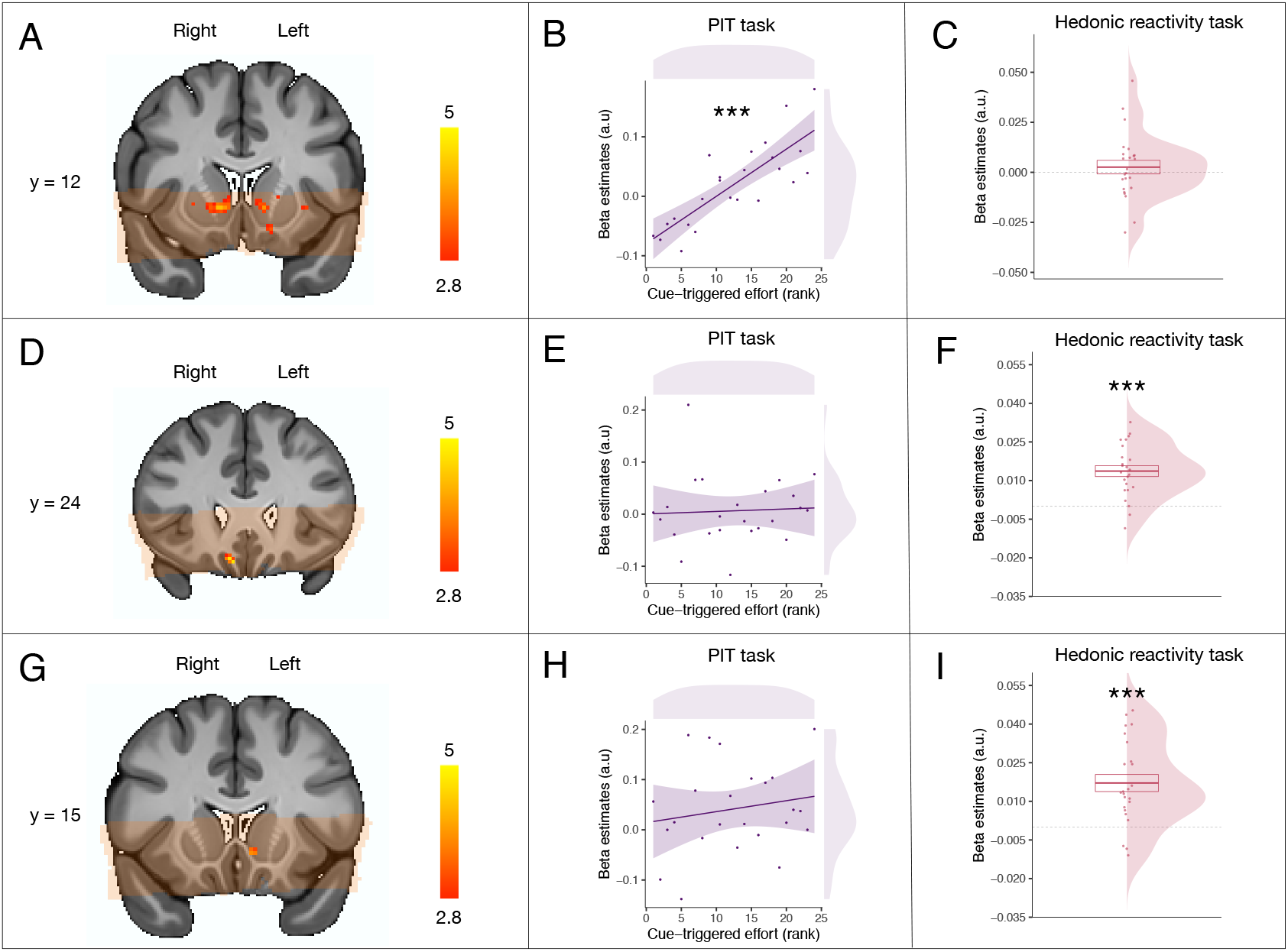
Neural correlates of Pavlovian-triggered motivation, and of sensory pleasure. **(A)** BOLD signal positively correlating with the magnitude of the PIT effect across participants in the ventral striatum. **(B)** Scatter plot showing the Pavlovian beta estimates (*CS*+ > *CS*−) extracted from the voxels within the ventral striatum correlating with the PIT effect against the strength of the behavioral PIT for each participant. **(C)** Overall mean across participants of the hedonic beta estimates (pleasure modulator) extracted from the voxels within the ventral striatum correlating with the PIT effect. **(D)** BOLD signal positively correlating with the magnitude of the hedonic pleasure experienced within participants in the mOFC. **(E)** Scatter plot showing the Pavlovian beta estimates (*CS*+ > *CS*−) extracted from the voxels within the mOFC correlating with the hedonic experience against the strength of the behavioral PIT for each participant. **(F)** Overall mean across participants of the hedonic beta estimates (pleasure modulator) extracted from the voxels within the mOFC correlating with the hedonic experience. **(G)** BOLD signal positively correlating with the magnitude of the hedonic pleasure experienced within participants within the ventral striatum. **(H)** Scatter plot showing the Pavlovian beta estimates (*CS*+ > *CS*−) extracted from the voxels within the ventral striatum correlating with the hedonic experience against the strength of the PIT effect for each participant. **(I)** Overall mean across participants of the hedonic beta estimates (*liking*) extracted from the voxels within the ventral striatum correlating with the hedonic experience. Orange overlay indicates from which brain areas the functional MRI data was acquired in all participants and were thus included in the statistical analysis. For display purposes, statistical *t*-maps are shown with a threshold of *p* < 0.005 uncorrected. Scale bar shows *t*-statistic. Error bars represent ±1 *SEM* adjusted for within participants designs. Asterisks indicate statistically significant differences.

To test whether these voxels were selectively activated for the PIT or whether they were also implicated in sensory pleasure, we extracted the beta estimates from these voxels during the hedonic reactivity task for our most sensitive pleasure contrast (see method). These beta estimates were not statistically different from 0 (*t*_(1,23)_ = 0.76, *p* = 0.45, *d*_*z*_ = 0.16, 95% CI = [−0.25, 0.57], *BF*_10_ = 0.002; see Fig. 5C).

#### Hedonic reactivity task

We report the results from our analyses within the predefined ROI in ventral striatum and mOFC using a height threshold of *p* < 0.005, with an extent threshold significant at *p* < 0.05 corrected for multiple comparisons.

Following the within-participants analysis typically used in hedonic reactivity tasks (*36*), we extracted the contrast correlating with the trial-by-trial experienced pleasantness reported by the participants. We found a statistically significant activation in the right mOFC (*k*^*thr*^ = 10; MNI [*xyz*] = [9 25 −18], *k* = 22, *β* = 0.013, 95% CI = [0.0093, 0.018], *SE* = 0.0021; see Fig. 5D,F). Moreover, we also found a significant activation in the left ventromedial subregion of the ventral striatum (*k*^*thr*^ = 15; MNI [*xyz*] = [−5 13 −5], *k* = 19, *β* = 0.017, 95% CI = [0.010, 0.024], *SE* = 0.0033; see Fig. 5D,F).

To test whether these voxels were selectively activated for sensory pleasure or whether they were also implicated in the PIT, we extracted the beta estimates (CS+ > CS−) from these clusters during the PIT task and correlated them with each participant’s PIT effect (increased effort during the CS+ compared to the CS−). This analysis did not reveal any statistically significant effect for the voxels in the mOFC (*F*_(1,22)_ = 0.06, *p* = 0.81, 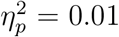, 90% CI = [0.00, 0.11], *BF*_10_ = 0.59; see Fig. 5E) or for the voxels in the ventral striatum (*F*_(1,22)_ = 0.69, *p* = 0.42, 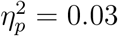, 90% CI = [0.00, 0.22], *BF*_10_ = 0.67; see Fig. 5H).

#### Comparison of ventral striatum subregions

To directly compare the activity of different subregions within the ventral striatum, we entered the activation of these regions in two separate statistical models. First, we ran a general linear model on the betas extracted during the PIT task (*CS*+ > *CS*−) testing the PIT effect on subregions of the ventral striatum (left ventromedial or bilateral dorsolateral) as function of the behavioral magnitude of the PIT effect across participants. This analysis revealed a statistically significant interaction between the sub-regions of the ventral striatum and the magnitude of the PIT effect (*F*_(1,22)_ = 6.58, *p* = 0.018, 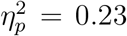, 90% CI = [0.03, 0.46], *BF*_10_ = 4.28), suggesting that the dorsolateral subregion of the ventral striatum was more involved in the PIT than the left ventromedial subregion of the ventral striatum. Second, we ran a general linear model on the betas extracted during the hedonic reactivity task testing the effect of the subregions of the ventral striatum (left ventro-medial or bilateral dorsolateral). This analysis revealed a main effect of ROI (*F*_(1,23)_ = 4.79, *p* = 0.039, 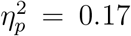, 90% CI = [0.01, 0.40], *BF*_10_ = 2.80), now suggesting that the left ventromedial subregion of the ventral striatum was more involved in the sensory pleasure than the dorsolateral subregion of the ventral striatum.

#### Pavlovian-triggered motivation and sensory pleasure within the core-like and shell-like divisions of the ventral striatum

To further test the differential contribution of the core and shell nuclei of the ventral striatum, we used the core-like and shell-like segmentation created by Cartmell et al., (*9*). We expected our general PIT effect to correlate with the activity of the core-like division and the sensory pleasure experience to correlate with the shell-like division.

First, we tested the implication of the core-like division in the PIT effect by extracting the beta estimates (CS+ > CS−) from within the core-like division during the PIT task and correlating them with the PIT effect of each participant (increased effort during the CS+ compared to the CS−). This analysis showed a statistically significant effect for the voxels in the core (*F*_(1,22)_ = 10.63, *p* = 0.004, 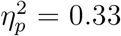, 90% CI = [0.08, 0.51], *BF*_10_ = 7.75; see Fig. 6B). To test whether the core-like division was selectively activated for the PIT effect or whether it was also implicated in sensory pleasure, we extracted the beta estimates from the core-like division during the hedonic reactivity task for our most sensitive pleasure contrast (see method). These beta estimates were not statistically different from 0 (*t*_(1,23)_ = 0.76, *p* = 0.35, *d*_*z*_ = 0.19, 95% CI = −0.21, 0.60], *BF*_10_ = 0.32).

**Fig. 6:**
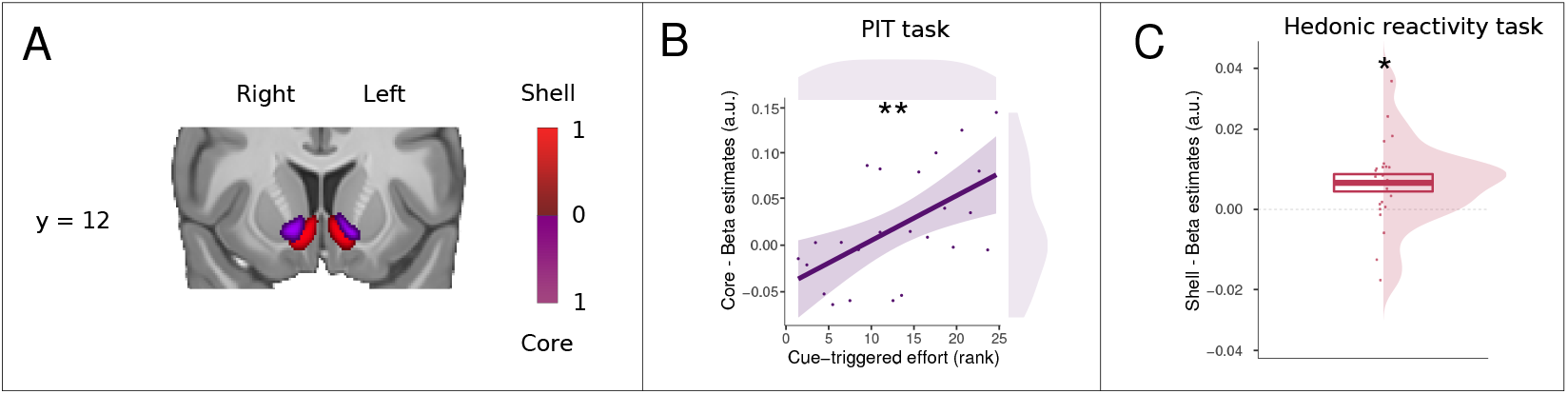
Pavlovian-triggered motivation and sensory pleasure within the core-like and shell-like divisions of the ventral striatum. **(A)** Probabilistic atlas from Cartmell and collaborators (*9*) depicting the core-like (in violet) and shell-like (in red) divisions of the human ventral striatum. The scale bar indicates the probability of the presence of a given division. **(B)** Scatter plot showing the Pavlovian beta estimates (*CS*+ > *CS*−) extracted from the core-like division of the ventral striatum against the strength of the PIT effect for each participant. **C**. Overall mean across participants of the hedonic beta estimates (*liking*) extracted from the shell-like division of the ventral striatum. *SEM* adjusted for within participants designs. Asterisks indicate statistically significant differences.

Second, we tested whether the shell-like division was involved in sensory pleasure by extracting the beta estimates from within the shell-like division during the hedonic reactivity task. These beta estimates were statistically different from 0 (*t*_(1,23)_ = 2.28, *p* = 0.032, *d*_*z*_ = 0.47, 95% CI = 0.03, 0.88], *BF*_10_ = 1.85; see Fig. 6C). To test whether the shell was selectively activated for sensory pleasure or whether it was also implicated in the PIT effect, we extracted the beta estimates (CS+ > CS−) from the shell-like division during the PIT task and correlated them with the PIT effect of each participant (increased effort during the CS+ compared to the CS−). This analysis did not reveal any statistically significant effect for the voxels in the shell-like division (*F*_(1,22)_ = 2.04, *p* = 0.17, 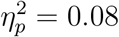, 90% CI = [0.00, 0.29], *BF*_10_ = 0.92).

## Discussion

This study investigated whether, like in rodents, different subregions of the human ventral striatum are differentially involved in the motivational and hedonic components of the affective processing of reward. With this aim, we combined a high-resolution fMRI protocol with a PIT task and a hedonic reactivity task using an olfactory reward, to try to maintain the paradigms as similar as possible to those used in animal research. This allowed us to measure Pavlovian-triggered motivation and the sensory pleasure experience for the same reward within the same participants. Our findings show evidence of dissociable contributions of different subregions of the ventral striatum to motivational and hedonic processes of reward. More specifically, we found that the dorsolateral subregion of the ventral striatum was more involved in Pavlovian-triggered motivation than in sensory pleasure, whereas the left ventromedial subregion of the striatum, similar to the mOFC, was conversely more involved in sensory pleasure than in Pavlovian-triggered motivation. Critically, when using the core-like and shell-like segmentation of the ventral striatum, our findings suggest that the Pavlovian-triggered motivation relied on the core-like division, whereas the sensory pleasure experience relied on the shell-like division of the ventral striatum.

Our study showing the involvement of the ventral striatum in Pavlovian-triggered motivation accords with findings from previous studies conducted in rodents (*10–12*) and humans (*15, 17, 18*). The nucleus accumbens has long been demonstrated to be implicated in PIT effects in rodents (*10–12*), with evidence of a dissociation between the shell and the core divisions as underlying two distinct forms of PIT: the outcome-specific and the general effects, respectively (*10*). In the outcome-specific PIT effects, a Pavlovian stimulus exerts a selective influence only invigorating a specific instrumental action associated with a specific reward, whereas in the general PIT effects, a Pavlovian stimulus triggers the invigoration of any instrumental responding irrespective of the specific reward the instrumental action is associated with (*37*). Whereas fMRI studies conducted in humans have found a correlation between the ventral striatum and the global PIT effects (*17, 18*), they have typically reported an activation of the dorsal striatum for outcome-specific effects (*27, 38*). The version of the PIT task we used in the present study did not allow us to distinguish between these two forms of Pavlovian influence on the instrumental action, nonetheless, the generic form of the PIT task we used is likely to reflect a general PIT effect, which is congruent with our selective activation of the core-like division of the ventral striatum. Importantly, the activation of the ventral striatum we found during our task replicates findings from a previous study using the same version of the task with a monetary reward (*15*). It is however important to note that there is a difference between our results and the results of the aforementioned study: Whereas Talmi and collaborators (*15*) found the ventral striatum to be correlated with the PIT effect within participants, we found the ventral striatum to be correlated with the magnitude of the PIT effect between participants. This difference could be driven by the behavior of our participants: Unlike Talmi et al. (*15*), we did not observe a strong effect of extinction during the PIT task. Therefore, our data showed less within-participant variability in terms of the PIT effect. By contrast, we observed a large variability in the magnitude of the PIT effect between participants, which provided the variance for the brain-behavior correlation analysis.

More generally, our findings highlighting the role of the more dorsolateral regions of the ventral striatum in reward motivation effects are congruent with prior work in the human fMRI literature showing that the involvement of the ventral striatum extends to the caudate in conditions with incentive actions and stimulus-driven motivational states (*2, 39*). Importantly, our findings further contribute to identifying the selectivity of the dorsolateral subregion of the ventral striatum in underlying the motivational component–as opposed to the hedonic component– of the affective processing of reward.

An important feature of our study is the use of an olfactory reward which triggers an immediate sensory pleasure experience, differently from other kinds of rewarding stimuli used in humans consisting in representations of rewards that will be delivered at a later stage (e.g., food pictures). This allowed us to specifically compare the involvement of distinct ventral striatum subregions during the PIT task and during the sensory pleasure experience triggered by the reward consumption, thus providing evidence for a functional dissociation. This methodological feature also provides a platform for a cross-species comparison between studies conducted in rodents and in humans. Our results are in line with findings from rodent studies showing that dopamine-agonist amphetamine injections within various subregions of the nucleus accumbens amplified the PIT effect, but not the hedonic response during reward consumption (*12, 40*). These studies have played a pivotal role in the formulation of the incentive salience hypothesis, which postulates that under some particular circumstances the motivational (i.e., wanting) and hedonic (i.e., liking) components of affective processing of the reward can be dissociated, thereby making organisms work for a reward that they will not necessarily appreciate once obtained–a key feature of compulsive reward-seeking behaviors such as addiction (*5*). However, in contrast to studies conducted in rodents, we were not able to determine in our study whether the observed activation of the ventral striatum dorsolateral subregion is related to dopaminergic activity. Interestingly, pharmacological studies conducted in humans (*41*) have shown that dopamine reduction decreases the magnitude of general appetitive PIT effects. Future studies might accordingly combine pharmacological manipulations with high-resolution fMRI protocols to shed more light on the neural mechanisms underlying the motivational and hedonic components of the affective processing of the reward. It is important to note there might be a caveat in the interpretation of our PIT effect in terms of Pavlovian influences. Similar to Talmi and collaborators (*15*), the Pavlovian learning task we used involved an instrumental component (i.e., pressing a key to discover whether the image was associated with the olfactory reward or not). However, the instrumental action had only limited predictive value in that the olfactory reward was delivered based on the CS image, and this was the case even when the instrumental action was not performed. Therefore, Pavlovian associative mechanisms were very likely to be dominant during Pavlovian learning task.

With respect to the hedonic component, we found the involvement of both a small subregion of the shell-like division of ventral striatum and the mOFC in the sensory pleasure experience during the reward consumption. The involvement of the mOFC in the sensory pleasure experience has long been established in human fMRI experiments (*20, 22*). By comparison, the involvement of a subregion of the ventral striatum in the sensory pleasure experience is more striking. fMRI studies conducted in humans have sometimes reported the involvement of the ventral striatum in hedonic reactions (*21,22*), but less consistently than the mOFC (*23*). Studies conducted in rodents have highlighted the presence of small “hedonic hotspots” in the shell division of the ventral striatum enhancing the hedonic experience when stimulated with opioids, orexin, or endocannabinoid (*13*). The combination of a high-resolution fMRI protocol and a hedonic reactivity task using an olfactory reward may have allowed us to detect the signal from such a small region in humans. Critically, in the hedonic reactivity task, we asked our participants to evaluate their hedonic experience during each trial. Our findings highlight the importance of this idiosyncratic measure, given that the perception of the pleasantness of the same odor varied in function of the habituation (i.e., whether the odor was presented twice or more in a row) and contrast effects (i.e., whether the pleasant odor was presented after an unpleasant odor). Both of these sequence effects are known to have a profound influence on chemosensory perception (*42–44*), hence the importance of taking into account the trial-by-trial variability within each participant.

Nevertheless, there are a number of factors that limit the comparisons that can be drawn between our findings and the findings from rodents studies that deserve to be discussed. First, our hedonic reactivity task consisted in explicit auto-reported hedonic evaluations rather the passive smelling with behavioral measures. It has been suggested that explicit evaluation tasks can have a different influence on hedonic activation compared to passive smelling tasks (*23*). Recently, it has been shown that the electromyographic signal from facial reactions during reward consumption can be successfully used as behavioral measure of pleasure without using auto-reports (*19*). Although we tried to implement such recordings in our study, the signal of these small facial movements was unfortunately not strong enough to be retrieved from the noise of the fMRI environment, we therefore did not have a behavioral measure of sensory pleasure in our hedonic reactivity task aside from auto-reports. Second, in our findings, we cannot determine whether the activation of the ventromedial subregion of the ventral striatum for sensory pleasure is modulated by opioids like in animals. Interestingly, a recent pharmacological study has shown that opioidergic manipulations through naltrexone led to a reduction of the implicit expression of sensory pleasure (*19*). Additional studies are thus necessary to assess whether opioidergic manipulations affect the involvement of the ventral striatum in sensory pleasure.

Notwithstanding these caveats, our results still provide evidence of dissociable contributions of the human ventral striatum subregions to the motivational and the hedonic component of the affective processing of the reward. These findings are important to further our understanding of the role of the ventral striatum in affective processes related to reward in both humans and other animals. A refined knowledge of these neural mechanisms might contribute to fostering novel insights into compulsive reward-seeking behaviors where motivational processes (such as wanting) are increased despite the absence of a related increase in hedonic processes (such as liking) (*5*). As Pavlovian influences have been proposed to play a pivotal role in a variety of psychiatric disorders and maladaptive behaviors, including addiction, binge eating, or gambling (*45–48*), modeling the interplay between Pavlovian incentive processes and hedonic processes could therefore have important implications for the understanding of psychological disorders.

## Materials and Methods

### Participants

We recruited 26 healthy participants at the University of Geneva. Participants were screened to exclude: (a) those with any previous history of neurological/psychiatric disorders, (b) those with any kind of olfactory disorder, and (c) those who were on a diet or seeking to lose weight. Moreover, participants were screened to include only those who perceived the chocolate odor used as a reward as pleasant. Data from two participants were excluded due to technical problems with their fMRI scans (one participant could not enter into the scanner because of a piercing and the images from the other participant could not be used because the table moved during the scanning session). We therefore used the data from the remaining 24 participants (11 females, age 26.56 ± 4.72 years). Participants were asked to fast for six hours prior to each experimental session. They gave their written informed consent and were paid 60 Swiss francs for their participation. The study protocol was approved by the Regional Research Ethics Committee in Geneva. The sample size was determined based on previous studies using similar high-resolution sequences on subcortical brain regions and similar tasks (*25–28*).

### Odor stimuli and presentation

The 12 olfactory stimuli (Aladinate, Cassis, Ghee, Indol, Leather, Paracresol, Pin, Pipol, Popcorn, Methyl salicylate, Yogurt, and Chocolate) were provided by Firmenich SA (Geneva, Switzerland).All odorants were diluted (20% v/v) in dipropylene glycol (DIPG), the control condition (odorless air) consisted of pure DIPG. The olfactory stimuli were selected based on pleasantness evaluations done in previous pilot studies on a visual analog scale going from 0 (“extremely unpleasant”) to 100 (“extremely pleasant”). The 11 neutral olfactory stimuli were selected based on their evaluations being more or less neutral (varying from *M* = 39 and *SD* = 21 to *M* = 66 and *SD* = 16) and the chocolate olfactory stimulus was selected to be used as the olfactory reward because it was consistently evaluated as being very pleasant (*M* = 82 and *SD* = 3).

The odors were delivered directly to the participants’ nostrils through a computer-controlled olfactometer with an air flow fixed at 1.5 L/min via a nasal cannula. There was a constant odorless air stream delivered throughout the experimental session, and the odorant molecules were delivered in this air stream without any change in the overall flow rate. The olfactory stimuli were thereby delivered rapidly and without thermal or tactile confounds, thus avoiding any change in the somatosensory stimulation (*49*).

### Mobilized effort

The mobilized effort was measured through an fMRI-compatible isometric handgrip (TDS121C) connected to the MP150 Biopac Systems (Santa Barbara, CA) with a 500 Hz sampling rate. The dynamic value of the signal was read by MATLAB (version 8.0) and used to provide participants with an online visual feedback (Psychtoolbox 3.0; for the visual interface implemented in MATLAB) that reflected the force exerted on the handgrip. This visual feedback was illustrated through the “mercury” of a thermometer-like image displayed on the left side of the screen (30 ° visual angle) that moved up and down according to the effort mobilized (see Fig. 1). The “mercury” of the thermometer-like display reached the top if the handgrip was squeezed with at least 50% or 70% (criterion varied every 1 s) of the participants’ maximal force. Note that we also recorded electromyographical activity from the zygomaticus and the corrugator muscles of the face of our participants. However, we could not systematically retrieve the signal of these recordings in the noise generated by the fMRI environment, and therefore did not include these results here. This data is nonetheless available with the rest of data presented here.

### Experimental Paradigm

The experiment consisted of two separate testing days. The first day was conducted outside the scanner, participants underwent the instrumental learning task, the Pavlovian learning task, and an odor selection task. The second day was conducted inside the scanner where the participants underwent the Pavlovian-Instrumental transfer test and the hedonic reactivity task.

#### Instrumental conditioning

Participants learned to squeeze a handgrip to trigger the release of the olfactory reward (same procedure as (*30*)). There were 24 trials (12 s) followed by an inter-trial interval (ITI; 4–12 s). During the trial, a fractal image (8° visual angle) and a thermometer were displayed in the center and on the left side of the screen, respectively. Participants were asked to squeeze the handgrip, thereby bringing the “mercury” of the thermometer up to the maximum and then down again, without paying attention to their squeezing speed. They were told that during the presentation of the thermometer display, there were three “special 1-s windows” and that if they happened to squeeze the handgrip during one of these time windows, they would trigger the release of the olfactory reward. They were also told that they were free to choose when to squeeze the handgrip and were encouraged to use their intuition. In reality, only two 1-s windows were randomly selected in each trial to be rewarded with the olfactory reward. If participants squeezed the handgrip with at least 50% or 70% (criterion varied every 1 s) of their maximal force during these time windows, a sniffing signal (a black asterisk; 2° visual angle) was displayed at the center of the fractal image and the olfactory reward was delivered. During the ITI, a fixation cross (2° visual angle) was displayed at the center of the screen and participants were asked to relax their hand to recalibrate their baseline force.

#### Pavlovian conditioning

Three initially neutral fractal images were attributed the Pavlovian roles of “baseline”, “CS+”, and “CS−”. The Pavlovian role of the fractal images was counter-balanced across participants. Each image was displayed at the center of the screen (visual angle of 8°). There were 36 trials (12 s) during which the CS+ or the CS− was displayed on the screen, followed by an ITI (12 s) during which the baseline image was displayed (procedure from (*30*)). During each trial, a target appeared every 4 s (on average) at the center of the CS image, three times per trial. Participants had to press the “A” key as fast as possible after they perceived the target, which was presented for a maximum of 1 s. Each time the CS+ image was displayed and the participant pressed the key, an olfactory reward was released; when the CS− image was displayed, odorless air was released. Participants were informed that the kind of odor released depended only on the CS image and not on the keypress task. In fact, the odor was released 1 s after the target onset when participants did not press the key during this interval. They were told about this aspect and it was moreover emphasized that the keypress task was a measure of their sustained attention and independent of the image-odor contingencies (see also (*15*)). During the ITI, the baseline image was displayed without any target, and no odor was released. After Pavlovian conditioning, participants evaluated the pleasantness of the images used as CS+, CS−, and baseline on a visual analog scale (from “extremely unpleasant” to “extremely pleasant”) presented at the center of the computer screen (visual angle of 23°). The order of the images was randomized across participants.

#### Odor selection task

Participants evaluated the pleasantness (from “extremely unpleasant” to “extremely pleasant”) and the intensity (from “not perceived” to “extremely strong”) of the 11 neutral odors, the olfactory reward, and the odorless air on visual analog scales displayed on a computer screen. Among the neutral odors, the odor rated as the most neutral (the closest to 50) and with the most similar intensity to the olfactory reward was selected to be used on the second day in the scanner for each participant.

#### PIT

The transfer test was administered on the second day, while participants were lying in the scanner. Participants were instructed to perform the same instrumental task as the day before, by squeezing the handgrip and keeping their gaze on the fractal image presented at the center of the screen. First, they completed three trials identical to those in instrumental conditioning (two “special 1-s” windows were rewarded), followed by six trials administered under partial extinction (one “special 1-s window” was rewarded). Immediately afterward, they performed the transfer test trials administered under extinction (no time window was rewarded). In the transfer test, the Pavlovian fractal images (CS+, CS−, or baseline) replaced the instrumental fractal image. The presentation order of the transfer test trials was randomized across the three stimuli (CS+, CS−, and baseline). There were five cycles of testing. In each cycle, each cue was presented three times consecutively, so that each Pavlovian stimuli was presented 15 times for a total of 45 transfer trials.

#### Hedonic reactivity task

The hedonic reactivity task was administered after the PIT, while participants were still lying in the scanner. Participants evaluated the pleasantness (from “extremely unpleasant” to “extremely pleasant”) and the intensity (from “not perceived” to “extremely strong”) of the three odor stimuli (rewarding, neutral, and odorless). Each odor release was preceded by a 3-s countdown, when the odor was released, the sniffing cue was presented at the center of the screen for 2.5 s. Afterward, the ratings were done on visual analog scales displayed on a computer screen and participants had to answer through a button-box placed in their hand. The answer to the question was self-paced, and the time participants took to answer was removed from the duration of the ITI (12s). There were 54 trials (18 per odor) consisting in six randomized cycles of presentation for each condition where the odor was administered three consecutive times per cycle.

### Behavioral analyses

Statistical analyses of the behavioral data were performed with R (*50*). For the analyses of variances (ANOVAs), we used the afex (*51*) and BayesFactor (*52*) packages. Adjustments of degrees of freedom using Greenhouse-Geisser correction were applied when the sphericity assumption was not met. We computed the Bayes factor (*BF*_10_) quantifying the likelihood of the data under the alternative hypothesis relative to the likelihood of the data under the null hypothesis using Bayesian ANOVAs (*53*). The Bayes factors reported for the main effects compared the model with the main effect in question versus the null model, while Bayes factors reported for the interaction effects compared the model including the interaction term to the model including all the other effects but the interaction term. Evidence in favor of the model of interest was considered anecdotal (1 < *BF*_10_ < 3), substantial (3 < *BF*_10_ < 10), strong (10 < *BF*_10_ < 30), very strong (30 < *BF*_10_ < 100) or decisive (*BF*_10_ > 100). Similarly, evidence in favor of the null model could also be qualified as anecdotal (0.33 < *BF*_10_ < 1), substantial (0.1 < *BF*_10_ < 0.33), strong (0.033 < *BF* 10 < 0.1), very strong (0.01 < *BF*_10_ < 0.033) or decisive (*BF*_10_ < 0.01) see (*54*). Partial eta squared 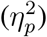 or Cohen’s *d*_*z*_ and their 90%*CI* or 95%*CI* are reported as estimates of effect sizes for the ANOVAs and the *t* tests, respectively.

### fMRI data acquisition

Functional imaging was performed at the Brain and Behavior Laboratory (University of Geneva) using a 3-Tesla MRI system (Magnetom Tim Trio, Siemens Medical Solutions) with a 32-channel receive array head coil for all the MR scanning sessions. The acquisition of the neuroimaging data was performed according to a high-resolution functional MRI sequence from Prévost et al. (*27*). We recorded 26 echo-planar imaging (EPI) slices per scan with a isotropic voxel size of 1.8-mm. Note that this volume is 4.6 times smaller than a fairly standard fMRI resolution with 3-mm isotropic voxels. Our scanner parameters were set at: echo time (TE) = 41 ms, repetition time (TR) = 2400 ms, field of view (FOV) = 180 × 180 × 39.6 mm, matrix size = 100 × 100 voxels, flip angle = 75°, no gap between slices. On account of our a priori regions of interest (ROIs), the acquired partial oblique axial T2^*^-weighted (T2^*^_*w*_) EPI mainly covered the striatum and the orbitofrontal cortex (OFC). The field of view was determined before the tasks and adjusted for each participant (see Fig. 4 and Fig. 5). We also acquired whole brain T1-weighted (T1_*w*_) images (isotropic voxel size = 1.0 mm), a whole brain reference functional image for the images’ co-registration, and dual-echo gradient *B*_0_ field maps to allow geometric correction of the EPI data.

### fMRI data preprocessing

We combined the Oxford Centre’s FMRIB (Functional Magnetic Resonance Imaging of the Brain) Software Library (FSL, version 4.1; (*55*)) with the Advanced Normalization Tools (ANTS, version 2.1; (*56*)) to tailor the preprocessing of high-resolution fMRI data for subcortical structures. FSL was used for brain extraction and realignment functional images. The functional images were automatically denoised using an independent components analysis and hierarchical fusion of classifiers (ICA-FIX). To achieve higher accuracy, the ICA-FIX classifier was trained on the present data set. Field maps were applied to correct geometric distortion (FUGUE). ANTS was used to diffeomorphically co-register the preprocessed functional and structural images to the California Institute of Technology (CIT168) brain template in the MNI space, using nearest-neighbor interpolation and leaving the functional images in their native 1.8-mm isometric resolution (*25*). Finally, we applied a spatial smoothing of 4-mm full width half maximum (FWHM).

### fMRI data analysis

The Statistical Parametric Mapping software (SPM, version 12; (*57*)) was used to perform a random-effects univariate analysis on the voxels of the image times series following a two-stage approach to partition model residuals to take into account within- and between-participant variance (*58, 59*). For the first-level, we specified a general linear model (GLM) for each participant. We used a high-pass filter cutoff of 1/128 Hz to eliminate possible low-frequency confounds (*15*). Each regressor of interest was derived from the onsets and duration of the stimuli and convoluted using a canonical hemodynamic response function (HRF) into the GLM to obtain weighted parameter estimates.

For each task, we created several GLMs with an increasing level of complexity and we performed model comparison and selection between the different GLMs using the MACS toolbox (Model Assessment, Comparison and Selection; (*60*)). We estimated cross-validated log model evidence (LME) for subject-level maps for each GLM. Performing cross-validated Bayesian model selection (BMS, (*61*)) on those maps allowed us to derive group-level exceedance probability (EP) maps. We selected the final model on the averaged EP across voxels within the striatum. The BMS procedure allowed us to select the best GLM for our group-level analysis given our data (see Fig. 6).

Group-level statistic *t*-maps were then created for each task by combining subject-level contrasts.

The multiple comparisons correction was done using the Analysis of Functional magnetic resonance NeuroImages software (AFNI; version 20.2.16 (*62*)). First, we used the 3dFWHMx function to estimate the intrinsic spatial smoothness of each dimension separately. Then, we used the new 3dClustSim function (*63*) to create–via Monte Carlo simulation to form those estimates–a cluster extent threshold corrected for multiple comparisons at *p* < 0.05 for a height threshold of *p* < 0.005 within the ROI. We report the extent threshold (*k*^*thr*^), the weighted parameter estimate (*β*) and the number of consecutive significant voxels at *p* < 0.005 within the cluster (*k*).

On account of our hypothesis, we used anatomical grey matter masks to define our a priori ROIs for cluster correction. We chose this method to remain faithful to the structural brain architecture. We used cytoarchitectural maps (*64*) to identify the mOFC, the Harvard-Oxford atlas to identify the ventral striatum, the cerebellum, as well as the thalamus, and the parcellation from Zhou et collaborators (*65*) for the olfactory cortex. We display non-masked statistical *t*-maps of our group results overlaid on a high-resolution template (CIT 168) in MNI space.

Finally, to further investigate the different involvement of the nuclei within the ventral striatum, we used the core-like and shell-like segmentation of the human ventral striatum created by Cartmell et al. (*9*) using a tractography-based approach. We used those probabilistic maps to test if the average activation within the core-like and shell-like divisions would map onto the motivational and hedonic components of the affective processing of the reward, respectively.

### Univariate test of Pavlovian-triggered motivation

We build four possible GLMs and used the BMS to select the one that was the most sensitive to variations in the striatum. The first GLM (*Between*) consisted of six regressors: (1) the onsets of reminder phase, (2) the onsets of the partial extinction phase, (3) the onsets of the PIT CS+, (4) the onsets of the PIT CS−, (5) the onsets of the PIT baseline, and (6) a parametric regressor of non-interest encompassing the phasic handgrip activity for each volume to account for residual movement. The second GLM (*Between+control*) was similar to the first one but we added a control regressor of non-interest to account for the repetition of the presentation of the same CS. The third GLM (*Within*) included the same regressors as the first GLM with an additional parametric modulator encompassing the force exerted on the handgrip during the presentation of the Pavlovian fractal images, whereas the fourth GLM (*Within+control*) additionally included the control regressor of non-interest to account for the repetition presentation of the same CS.

Results of the BMS showed that the second GLM had the best fit within the striatum (see Fig. 7).

**Fig. 7:**
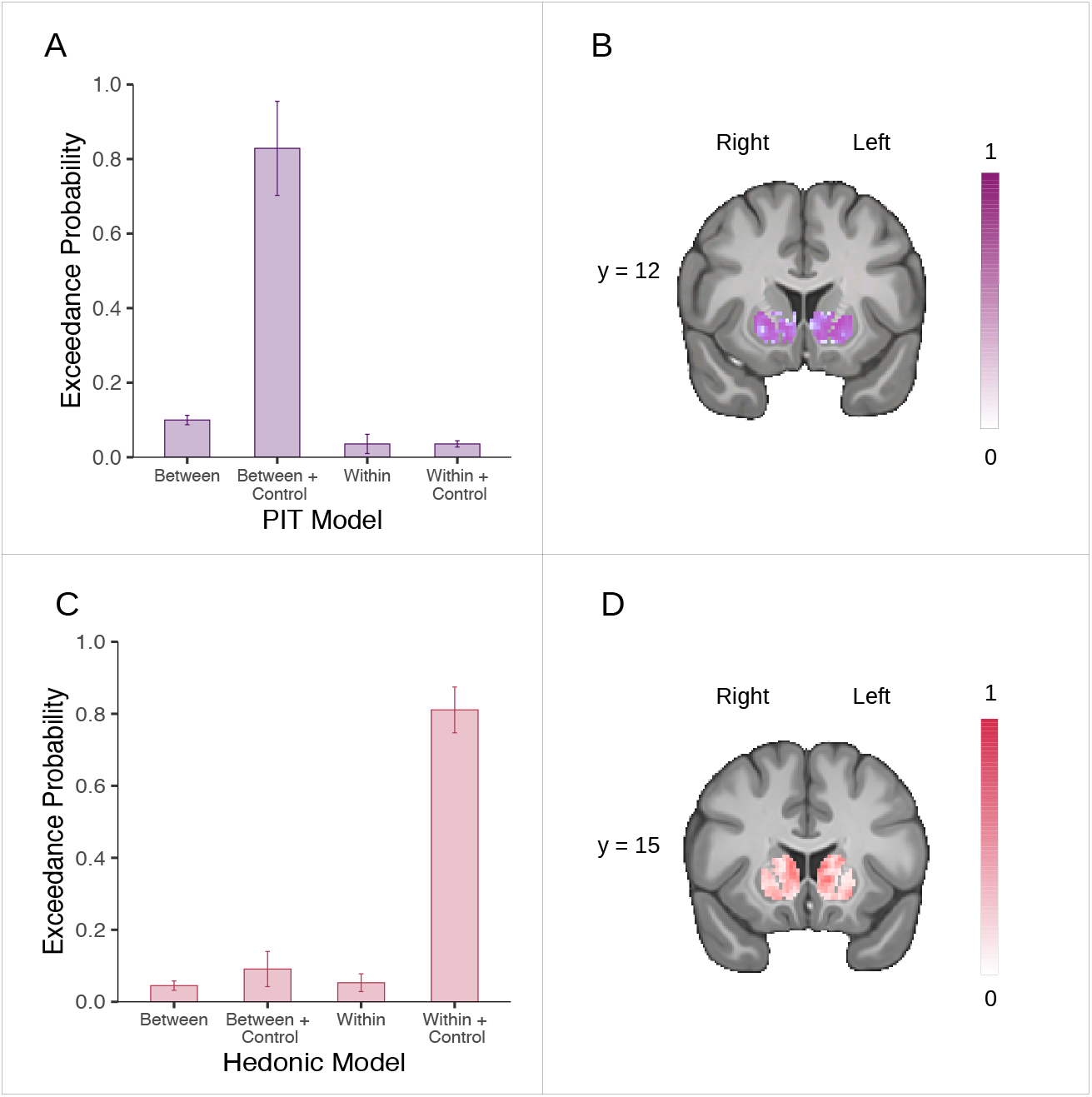
Voxel-wise Bayesian model selection in the striatum. **(A)** Mean exceedance probability across voxels within the striatum for the PIT models. **(B)** Likeliest frequency map of the winning PIT model (Between + Control). Mean exceedance probability across voxels within the striatum for the hedonic models. **(D)** Likeliest frequency map of the winning hedonic model (Within + Control). Scale bars show the proportion of subjects in which the winning model is the optimal model. Error bars represent ±1 *SD*.

The main group-level contrast was derived from the linear difference between the CS+ and the CS− conditions correlated with the behavioral PIT effect. The behavioral PIT effect was computed by taking each participant’s average number of squeezes exerted during the CS+ subtracted by the average in the CS− condition, which we then de-meaned (*66*) and rank-transformed to normalize the distribution.

Finally, a control GLM was also computed with the onsets of every single squeeze during the whole task independently of the experimental condition. This aimed to validate our task and to control the quality of the BOLD signal by verifying whether the main effect of squeezing frequency activated motor regions included in our field of view.

### Univariate test of the pleasure experience

We built four possible GLMs and used the BMS to select the one that was the most sensitive to variations in the striatum. The first GLM (*Between*) consisted of six regressors: (1) the onsets of the trial, (2) the onsets of the reception of the pleasant odor, (3) the onsets of the reception of the neutral odor, (4) the onsets of the reception of the odorless air, (5) the onsets of the question about odor pleasantness, and (6) the onsets of the question about odor intensity.

The second GLM (*Between+control*) was similar to the first one but we added a control regressor of non-interest to account for the repetition of the presentation of the same odor.

The third GLM (*Within*) consisted of seven regressors: (1) the onsets of the trial, (2) the onsets of the reception of an odor modulated by (3) the trial-by-trial ratings of the perceived pleasantness and (4) the trial-by-trial ratings of the perceived intensity, (5) the onsets of the reception of the odorless air, (6) the onsets of the question about odor pleasantness, (7) the onsets of question about odor intensity. The two modulators locked on the onset of the reception of the odors were competing for variance, so that they would each represent their individual explained variance (*66*).

The fourth GLM (*Within+control*) was identical to the second one but we added two additional regressors of non-interest accounting for the repetition in the presentation of the same odor and whether the odor presented at a given trial was more or less pleasant than the preceding trial.

Results of the BMS showed that the third GLM had the best fit within the striatum (see Fig. 7).

The main group-level contrast was derived from the parametric modulation of pleasantness on the odor reception.

Finally, a control GLM was also computed with the onset of the reception of the odors and the onset of the odorless air reception, as well as the perceived intensity of the odors as a second-level modulator. This aimed at validating our task and quality control of the BOLD signal by verifying whether the main effect of odor activated the olfactory regions in our field of view.

## General

The authors would like to thank Dr. Leonardo Ceravolo for the discussion on the analysis and Dr. Vanessa Sennwald for her insightful comments on the manuscript. The authors would also like to acknowledge the precious contribution and support at the early stages of this project of our colleague and friend Dr. Charlotte Prévost who passed away in March of 2016.

## Author contributions

D.S., S.D., and E.R.P. designed the experimental task. D.S., P.V., S.D., and E.R.P. discussed the experimental design. D.C. took part in the choice and preparation of the odors and the settings of the olfactometer. Y.S. and E.R.P. collected the data. D.M.T, S.D., and E.R.P. analyzed the data. D.M.T and E.R.P. wrote the manuscript. All authors discussed the results and commented on the manuscript at all stages.

## Data and materials availability

Raw, de-identified MRI data are available at the Open Neuro platform [https://openneuro.org/datasets/ds003487/]. Computer code used for preprocessing and analyzing the data is available in a publicly hosted software repository [https://github.com/evapool/VS_AffectiveResponse].

